# Detection of Cardiac O-GlcNAcylation via Subcellular Fractionation and Dual Antibody Analysis in Pressure Overload Cardiac Hypertrophy

**DOI:** 10.1101/2025.10.14.682429

**Authors:** Dolena Ledee, Wei Zhong Zhu, Aaron K Olson

## Abstract

Protein O-GlcNAcylation is a dynamic post-translational modification with emerging roles in cardiac pathophysiology. The availability of differenct pan-specific antibodies to assess global O-GlcNAc levels, and variability in Western blot results has hindered cross-study reproducibility and interpretation. In this study, we applied optimized immunoblotting protocols using both CTD110.6 and RL2 O-GlcNAc antibodies, alongside subcellular fractionation, to investigate temporal and sex-specific changes in cardiac O-GlcNAcylation during pressure overload hypertrophy (POH) from transverse aortic constriction (TAC) during early (1-week POH, 1wTAC) and chonic (6-weeks POH, 6wTAC) POH in mice. Global O-GlcNAc levels were elevated in early POH and returned to baseline in chronic POH, consistent across both antibodies and sexes. Subcellular fractionation revealed persistent O-GlcNAc elevations in cytoplasmic and membrane fractions in chronic POH for both sexes, which were not detected in unfractionated samples. Female mice exhibited significantly higher O-GlcNAc levels than males during POH, particularly at early POH, highlighting sex-specific regulation. OGT and OGA protein levels also varied by compartment and sex, suggesting differential enzymatic control. In conclusion, our findings underscore the importance of methodological rigor in O-GlcNAc detection and demonstrate that fractionation enhances sensitivity to subtle changes in cardiac O-GlcNAcylation. Our principal new findings are protein O-GlcNAcylation dysregulation continues from early POH (1wTAC) into chronic POH (6wTAC groups) along with showing differences in O-GlcNAc levels between males and females during POH. These results provide new insights into the temporal and sex-dependent dynamics of O-GlcNAc signaling in POH and support its potential as a therapeutic target in cardiovascular disease.

## INTRODUCTIOIN

*O*-GlcNacylation is the reversible post-translational modification involving the addition of *O*-linked b-*N*-acetylglucosamine (*O*-GlcNAc) to serine/threonine residues on proteins. Despite being regulated by only two enzymes, OGA ( *O*-GlcNAcase) for removal and OGT (*O*-GlcNAc transferase) for addition, this modification dynamically affects hundreds to thousands of proteins influencing their activity, stability, and subcellular localization [1]. Dysregulation of global *O*-GlcNAc has been implicated in various diseases, including cardiac pathologies [2–4]. Our lab and others have observed elevated global O-GlcNAc levels in models of cardiac hypertrophy [2,5,6]. However, the functional consequences from this change are complex, as higher *O*-GlcNAc levels can exert both cardioprotective or detrimental effects depending on the context [4,7,8]. Adding to this complexity, *O*-GlcNAc dynamics demonstrate temporal and gender-specific patterns [3]. For instance, although male and female mice show similar chronological changes in cardiac O-GlcNAc levels following pressure overload hypertrophy (POH), the expression of OGT and OGA differs between sexes [3]. Additionally, we found thyroid hormone perturbations in aged mice can differentially alter global cardiac O-GlcNAc levels between sexes [9]. Together, these findings emphasize the nuanced regulation of cardiac O-GlcNAcylation and suggest that its context-dependent effects may be shaped by temporal, and sex-specific factors, warranting further investigation into its role in cardiovascular disease.

An ongoing challenge in the O-GlcNAc field is the variability observed in published western blot data [10]. Due to the limited availability of site-specific O-GlcNAc antibodies, most studies rely on pan-specific antibodies to semi-quantitatively assess global O-GlcNAc levels. However, each pan-specific antibody recognizes distinct epitopes and results in different banding patterns [11]. Notably, even when the same antibody is used, western blot results can vary significantly between and even within laboratories, complicating data interpretation and reproducibility [10,11]. This variability poses a significant barrier to reproducibility and cross-study comparisons, underscoring the need for standardized approaches in O-GlcNAc research.

Recent efforts have aimed to address this issue by optimizing immunoblotting methods for cardiac tissue. Zou et al. developed conditions for use with the CTD110.6 antibody for detection of a wider range of O-GlcNAcylated proteins-ranging from < 30 to around 450 kD [10]. Separately, Narayanan et al. refined protocols using the RL2 antibody, which demonstrated superior reactivity toward cardiac contractile proteins compared to CTD110.6 [11].

We previously found global cardiac O-GlcNAc levels increase in early POH (1-week post-transverse aortic constriction or TAC) and fall to Sham levels with chronic POH (6-weeks post-TAC), but that study used the RL2 antibody only and without optimized conditions [3]. In the current study, our goal was to determine whether applying improved methods and using both the RL2 and CTD110.6 antibodies provides new understanding on the temporal changes in O-GlcNAc levels from POH. We further tested whether performing O-GlcNAc immunoblots on cellular compartments using fractionation provides additional information on dynamic O-GlcNAc changes over the standard, non-fractionated approach. Finally, given the sex-differences in cardiac O-GlcNAc levels, we performed our studies in both males and females.

## METHODS

### Ethics Statement

This investigation conforms to the *Guide for the Care and Use of Laboratory Animals* published by the National Institutes of Health (NIH publication No. 85-23, revised 1996) and were reviewed and approved by the Office of Animal Care at Seattle Children’s Research Institute. All materials used in this study are available commercially from the indicated vendors.

### Mice

Pressure overload affects cardiac function and hypertrophic growth differently in males and females, so we performed these experiments in a both genders [12,13]. Additionally, our prior work showed that males and females can have differential regulation of cardiac O-GlcNAcylation [3]. C57Bl/6j mice were purchased from the Jackson Laboratory (Bar Harbor, ME). All mice were allowed free access to water and Teklad #7964 chow (Envigo, East Millstone, NJ, USA). Teklad ¼ inch corncob bedding was utilized in the cages (Envigo, East Millstone, NJ, USA). The mice were on a 12-hour light cycle. Mice were randomly assigned to their experimental groups.

### Experimental set-up

Our goal was to re-assess the temporal changes in O-GlcNAc levels during POH utilizing recently described optimized techniques for evaluating global levels as well as fractionated cellular compartments. We used TAC surgery to create POH. We utilized the following experimental groups which are the same as our prior study: early POH at 1-week after TAC (1wTAC), chronic POH at 6-weeks after TAC (6wTAC), and Sham. We previously found sham for 1-week or 6-weeks did not affect global O-GlcNAc levels or key O-GlcNAc regulatory proteins, so we used a single Sham group composed of approximately equal numbers of mice from 1-and 6-weeks post-Sham surgery [3].

### Surgery

We performed TAC or Sham surgery as previously described in our lab [3,4]. Mice were between the ages of 14-15 weeks at surgery. Briefly, mice were initially anesthetized with 3.5% isoflurane in 100% O2 at a flow of 1 LPM and then maintained with 2.5% isoflurane for the duration of the surgery. A midline sternotomy was performed to expose the aorta. To get a similar pressure gradients between the sexes, we constricted the transverse aorta with 7–0 silk suture tied against a 22-gauge blunt needle for males and 23-gauge for females. Sham mice underwent similar surgery but did not have the suture tightened around the aorta. For pain control, mice received intraperitoneal buprenorphine extended release (1.95–3.25 mg/kg IP) pre-operatively which provides analgesia for 72 hours. To maintain a consistent degree of pressure overload, we excluded any mice from analysis with an aortic constriction pulse wave Doppler velocity < 3.5 m/sec.

### Echocardiograms

We performed echocardiograms as previously described in our lab [14–16] to measure cardiac function and the degree of aortic constriction from the surgery. Briefly, mice were initially sedated with 3% isoflurane in O2 at a flow of 1 LPM and placed in a supine position at which time the isoflurane is reduced to 1.0-1.5% administered via a small nose cone. ECG leads were placed for simultaneous ECG monitoring during image acquisition. Mice were maintained at a temperature of greater than 37.0° C throughout the echocardiogram. Echocardiographic images were performed with a Vevo 3100 machine using a MX550D or MX250 transducers (VisualSonics, Inc, Toronto, Canada). M-Mode measurements at the midpapillary level of the left ventricle (LV) were performed at end-diastole (EDD) and end-systole (ESD) to determine LV function via the fractional shortening [(LVEDD-LVESD)/LVEDD * 100] in a parasternal short axis mode for at least three heart beats. We measured the pulse wave Doppler velocity across the aortic constriction site for a non-invasive estimate of the peak instantaneous pressure gradient (PIPG) across the construction site as calculated by the simplified Bernoulli equation (PIPG = 4*velocity^2^). The echocardiogram reader was blinded to the treatment. We previously found that isoflurane anesthesia during echocardiograms increases myocardial total protein O-GlcNAc levels for several hours after the procedure (unpublished data), so the echocardiograms were always performed one day prior to sacrifice.

### Whole cell protein isolation

The protocol for isolating whole cell protein from mouse cardiac tissue was previously described by our lab [3,4,6]. Briefly, protein was isolated with a RIPA lysis buffer (catalog #: 20-188, Millipore Sigma, Burlington, MA, USA) containing phosphatase inhibitor (Thermo Fisher Scientific, Waltham, MA, USA) plus the OGA inhibitors PUGNAc 20 μM (Toronto Research Chemical, North York, ON, Canada) and Thiamet-G 1μM (Adooq Bioscience, Irvine, CA, USA). Twenty-five micrograms of total protein extract from mouse heart tissue was separated by electrophoresis and transferred to PVDF membrane.

### Sequential subcellular fractionation method

We performed this method based on protocols from Baghirova et. al. [17] and Zou et. al. [10] whereby increasing detergent strength of lysis buffer releases protein from different cell compartments.

### Cytoplasmic protein fraction

∼20 mg of heart tissue was dounce homogenized on ice, 20 counts, in cold lysis buffer A: 50 mM Tris-HCl, 5 mM EGTA, 2 mM EDTA, 5 mM DTT, 0.05% digitonin plus 1X protease inhibitors. Samples were placed on ice for 30 minutes, and vortexed every 5 minutes. The samples were then centrifuged at 4° C for 10 minutes at 14,000 rpm and resulting supernatant transferred to a precooled tube.

### Membrane enclosed protein fraction

The pellet from the cytoplasmic protein fractionation was washed twice in cold lysis buffer A and centrifuged at 4° C for 10 minutes at 14,000 rpm. The supernatant from each was discarded. The pellet was then resuspended in cold lysis buffer B: 50 mM Tris-HCl, 5 mM EGTA, 2 mM EDTA, 5 mM DTT, 0.05% digitonin, 1% Triton X-100 plus 1X protease inhibitors. Samples were placed on ice for 30 minutes, and vortexed every 5 minutes. The samples were then centrifuged at 4° C for 10 minutes at 14,000 rpm and resulting supernatant transferred to a precooled tube.

### Soluble nuclear protein fraction

The pellet from the membrane enclosed protein fractionation was washed twice in cold lysis buffer B and centrifuged at 4° C for 10 minutes at 14,000 rpm. The supernatant from each was discarded. The pellet was then resuspended in cold lysis buffer C: 20 mM Hepes, pH 8, 2 mM EDTA, 0.5 mM DTT, 25% glycerol, 1.5 mM MgCl_2_, 300 mM KCl, 0.2 mM PMSF, 1U/ml benzonase (catalog #70746, Millipore Sigma, Burlington, MA) plus 1X protease inhibitors. Samples were placed on ice for 40 minutes, and vortexed every 10 minutes. The samples were then centrifuged at 4° C for 10 minutes at 14,000 rpm and resulting supernatant transferred to a precooled tube. Samples were desalted prior to PAGE using the Zeba^TM^ spin desalting columns, 7K MWCO (catalog # 89882, Thermo Fisher Scientific, Bothell, WA) per manufacturer’s instructions.

### Non-ionic detergent insoluble protein fraction (Myofilament and contractile proteins)

The pellet from the nuclear protein fractionation was washed twice in cold lysis buffer C and centrifuged at 4° C for 10 minutes at 14,000 rpm. The supernatant from each was discarded. The pellet was then resuspended in lysis buffer D: 50 mM Tris-HCl, 1% SDS, 5% glycerol, 5 mM DTT plus 1X protease inhibitors. Samples incubated at 20° C for 10 minutes, and vortexed every 3 minutes. The samples were then centrifuged at 20° C for 10 minutes at 14,000 rpm and resulting supernatant transferred to a fresh tube. All samples were stored at -80° C until subsequent analysis.

### Antibodies

We used the following primary antibodies: From Santa Cruz Biotechnology, Dallas, TX: GAPDH (catalog # sc-25778, assess cytoplasmic fraction), WTAP (catalog # sc-55438, assess nuclear fraction), SERCA (catalog # sc-376235, assess membrane enclosed fraction), Troponin C (catalog # sc-20642, assess myofilament/contractile fraction). From Proteintech Group, Rosemont, IL: ATP5C1 (catalog # 10910-1-AP, assess membrane enclosed fraction), OGA (catalog # 14711-1-P), OGT (catalog # 11576-2-AP), BAML1 (catalog # 14268-1-AP). From Cell Signaling, Danvers, MA: O-GlcNAc CTD110.6 (catalog # 9875), H2AX (catalog # 2595, asses insoluble protein fraction). From Novus Biologicals, Centennial, CO: O-GlcNAc RL2 (catalog # NB300-524).

### Western Blots

To confirm fractionation efficiency, 20 µg of lysate from each fraction was run on a 4-20% gradient gel (catalog # 5671095, BioRad Laboratories, Hercules, CA) and transferred to Immun-Blot® PVDF membrane (catalog # 1620177, BioRad Laboratories, Hercules, CA). Membranes were cut horizontally as to assess differentially sized proteins targeting different fractions on the same blot using the antibodies listed: GAPDH, WTAP, SERCA, ATP5C1, H2AX, and Troponin C.

To assess O-GlcNAc levels, 25 µg of whole cell lysate, 5 µg of lysate from the nuclear fraction, or 20 µg of lysate from the cytoplasmic, membrane enclosed, or insoluble pellet fractions were run on a 7.5% gradient gel (catalog # 5678025, BioRad Laboratories, Hercules, CA) and transferred to Immun-Blot PVDF membrane. Transfer of proteins to the PVDF membranes utilized a discontinuous Tris-CAPS buffer for semi-dry transfer. The anode buffer included 60 mM Tris, 40 mM CAPS, pH 9.6, plus 15% methanol, whereas the cathode buffer included 60 mM Tris, 40 mM CAPS, pH 9.6, plus 0.1% SDS. The membranes were probed initially with: O-GlcNAc CTD110.6, O-GlcNAc RL2, and OGT. The membranes containing fractionated proteins were subsequently stripped by washing 2 times, 15 minutes each, with 100 mM dithiothreitol, 2% (wt/vol) SDS, 62.5 mM Tris·HCl, pH 6.7, at 70℃, followed by three 10-min washes with TBST for additional antibody analysis with OGA. The membranes containing whole cell lysate were stripped for 15 minutes at 60℃ with Restore^TM^ PLUS Western Blot stripping buffer (catalog #46430, Thermo Fisher Scientific, Bothell, WA) followed by three 10-min washes with TBST for additional antibody specificity analysis with the RL2 or CTD110.6 antibody in the presence of 100 mM GlcNAc (catalog # A8625, Millipore Sigma, Burlington, MA). Immunoblots were normalized to total protein staining by Pierce Reversible Protein Stain Kit for PVDF Membranes (catalog # 24585, Thermo Fisher Scientific, Bothell WA), which are shown in the figures. Nuclear fraction immunoblots were normalized to strongest band after staining with the SYPRO^TM^ Ruby protein blot stain (catalog # S11791, Thermo Fisher Scientific, Bothell WA) or the transcriptional activator, BMAL1. Immunoblots and protein staining were measured by densitometry analysis using ImageJ software (National Institute of Health).

Membranes incubated with RL2, OGT and OGA antibodies used 5% milk for blocking, primary and secondary antibody incubation. The membranes incubated with CTD antibody used 3% BSA (#A7906,Thermo Fisher Scientific, Rockford, IL, USA).

Due to the RL2 antibody being raised in mice and the secondary anti-mouse antibody can react with IgG from the tissue resulting in a non-specific band at either 55 kD (from IgG heavy chain) or 25 kD (from IgG light chain), we used a light chain specific secondary antibody (catalog #: 115-035-174 Peroxidase AffiniPure goat anti-mouse IgG, light chain specific; Jackson ImmunoResearch Laboratories, Inc., West Grove, PA) for our RL2 immunoblots to allow for unobstructed detection of RL2 O-GlcNAc bands in the 55 kD region, and excluded the 25 kD band during quantification. Total (global) O-GlcNAc levels for both antibodies were measured by densitometry of the entire western blot lane and normalized to the total protein staining by Thermo Fisher Scientific Pierce Reversible Protein Stain Kit for PVDF Membranes (catalog #: 24585, Thermo Fisher Scientific, Rockford, IL, USA). If the event that global unfractionated O-GlcNAc levels were similar between groups, we also assessed for individual band differences.

### Data visualization and statistical analysis

All reported values are mean ± standard error of the mean (SEM). We compared experimental groups by pairwise t-test with Bonferroni correction for multiple groups when applicable. Criterion for significance was *p* < 0.05 for two-groups (example male-female comparisons) and p < 0.017 when comparing three groups. We preformed data visualization using GraphPad Prism version 8.3.3 (GraphPad Software).

## RESULTS

### Echocardiographic and Morphometric Assessment of Pressure Overload Hypertrophy (POH)

We used echocardiography and morphometric analyses to characterize the cardiac response to pressure overload in our experimental model (Figure 1). We quantified aortic constriction severity by PIPG across the constricted site by echocardiogram. As expected, male mice exhibited significantly elevated PIPG ( p<0.017) in both 1wTAC (55.9 ± 2.8 mmHg) and 6-week TAC (51.9 ± 1.3 mmHg) compared to Sham controls (mean 2.0 ± 0.1 mmHg). Female mice also showed significantly increased PIPG ( p<0.017) in 1wTAC (54.3 ± 1.6 mmHg) and 6wTAC (63.1 ± 3.7 mmHg) versus Sham (2.4 ± 0.1 mmHg).

**Figure 1.**
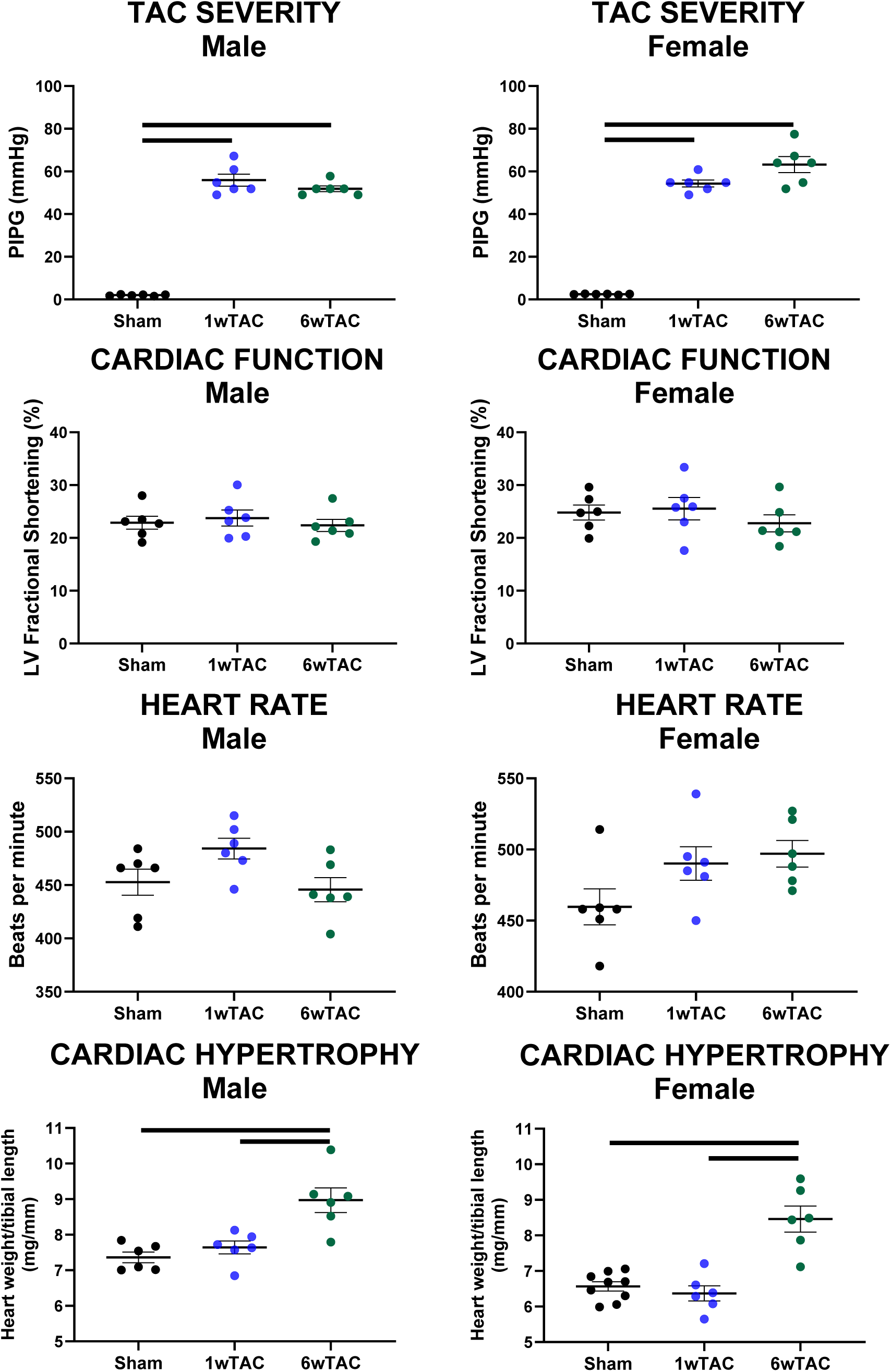
Echocardiographic and morphometric measurements. Heart rate measured during echocardiogram. All quantitative data are mean ± S.E.M (*n* ≥ 6); Statistical significance was based on unpaired *t*-test with a Bonferroni correction (*p* < 0.017). Brackets denote significance between indicated groups. TAC, transverse aortic constriction; LV, left ventricle.

Left ventricular systolic function, assessed by fractional shortening, remained preserved across all groups and sexes. Heart rate during echocardiography was also unaffected by TAC. Cardiac hypertrophy, evaluated using the ratio of biventricular heart weight to tibial length, was present in only the 6wTAC groups for both sexes ( p<0.017).

### Western Blot Analysis: Global O-GlcNAc Levels

We first evaluated for temporal changes in global O-GlcNAc levels during POH separately in male and female mice using both antibodies (Figures 2). In both sexes, global O-GlcNAc levels were significantly elevated at 1wTAC compared to Sham and returned to baseline Sham levels by 6wTAC. These findings were consistent across both antibodies and aligned with our previous study using RL2 alone [3]. Because global O-GlcNAc levels were similar between Sham and 6wTAC, we assessed for differences on individual bands and found only one significant band difference for the female RL2 group at approximately 102 kD (p = 0.014).

**Figure 2.**
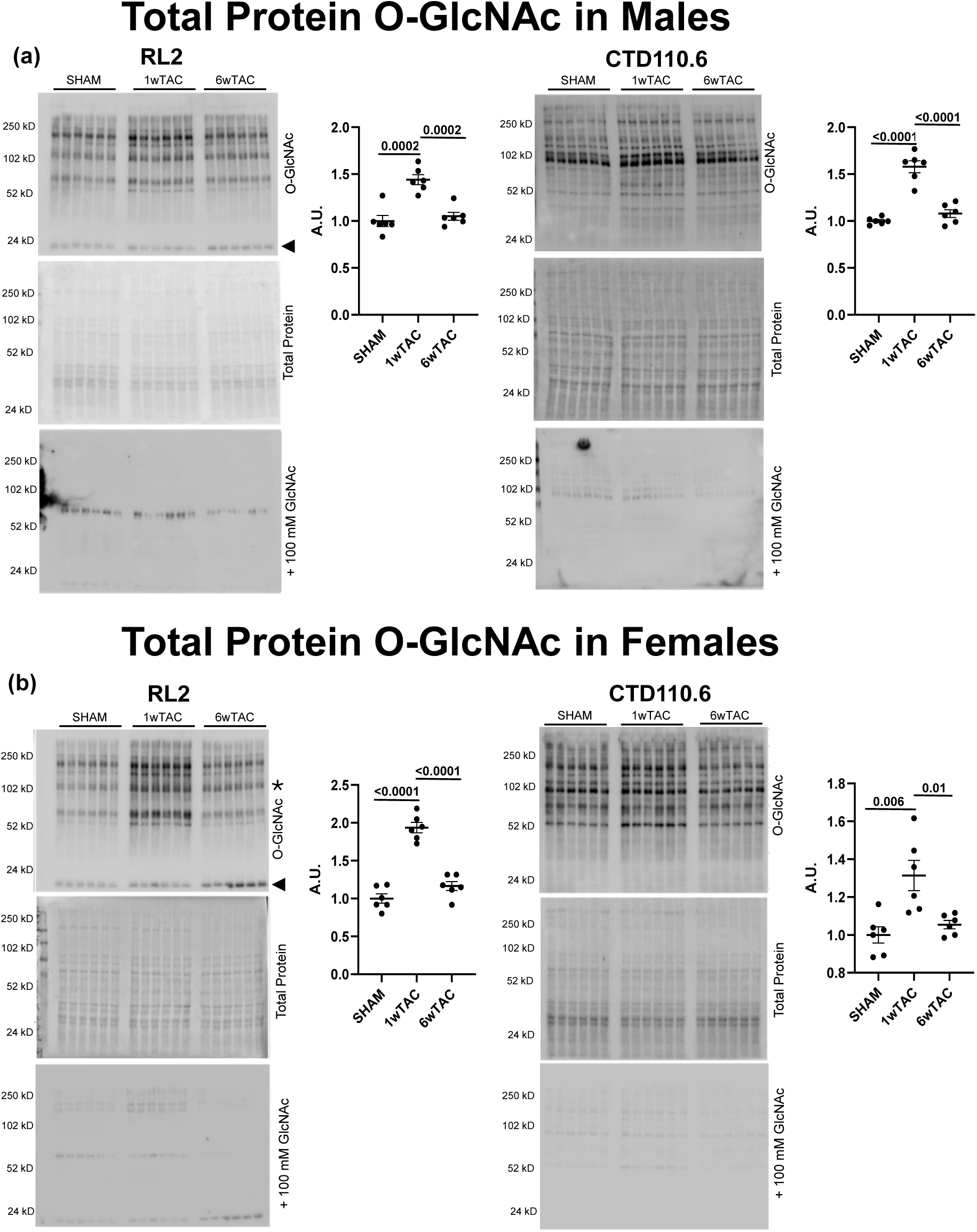
Global unfractionated cardiac protein O-GlcNAc levels detected by RL2 or CTD110.6 antibodies. Males (a) and females (b) comparing Sham, 1wTAC and 6wTAC. All values are arbitrary units (A.U.) normalized to total protein levels. Statistical significance was based on unpaired *t*-test with a Bonferroni correction (*p* < 0.017). Brackets denote significance between indicated groups. 3, κ (kappa) light chains IgG. *, significant band difference between the sham and 6wTAC (p = 0.014).

The RL2 and CTD110.6 antibody specificity was assessed in the presence of the blocking agent GlcNAc. We found a lack of O-GlcNAc bands which confirms the specificity of our O-GlcNAc western blots (Figure 2). Please note that the exposure time for the competitive GlcNAc western blots was longer (around 4 minutes) than the standard O-GlcNAc blots (typically 15 seconds).

We then compared O-GlcNAc levels between sexes for each experimental group. There were no differences in global O-GlcNAc levels for Sham hearts with either antibody. In the 1wTAC group, female mice exhibited significantly higher global O-GlcNAc levels than males using both CTD110.6 (p = 0.045) and RL2 (p = 0.03) antibodies (Figure 3). At 6wTAC, sex differences varied by antibody with RL2 showing increased global O-GlcNAc in females (p = 0.003), while CTD110.6 showing no sex differences.

**Figure 3.**
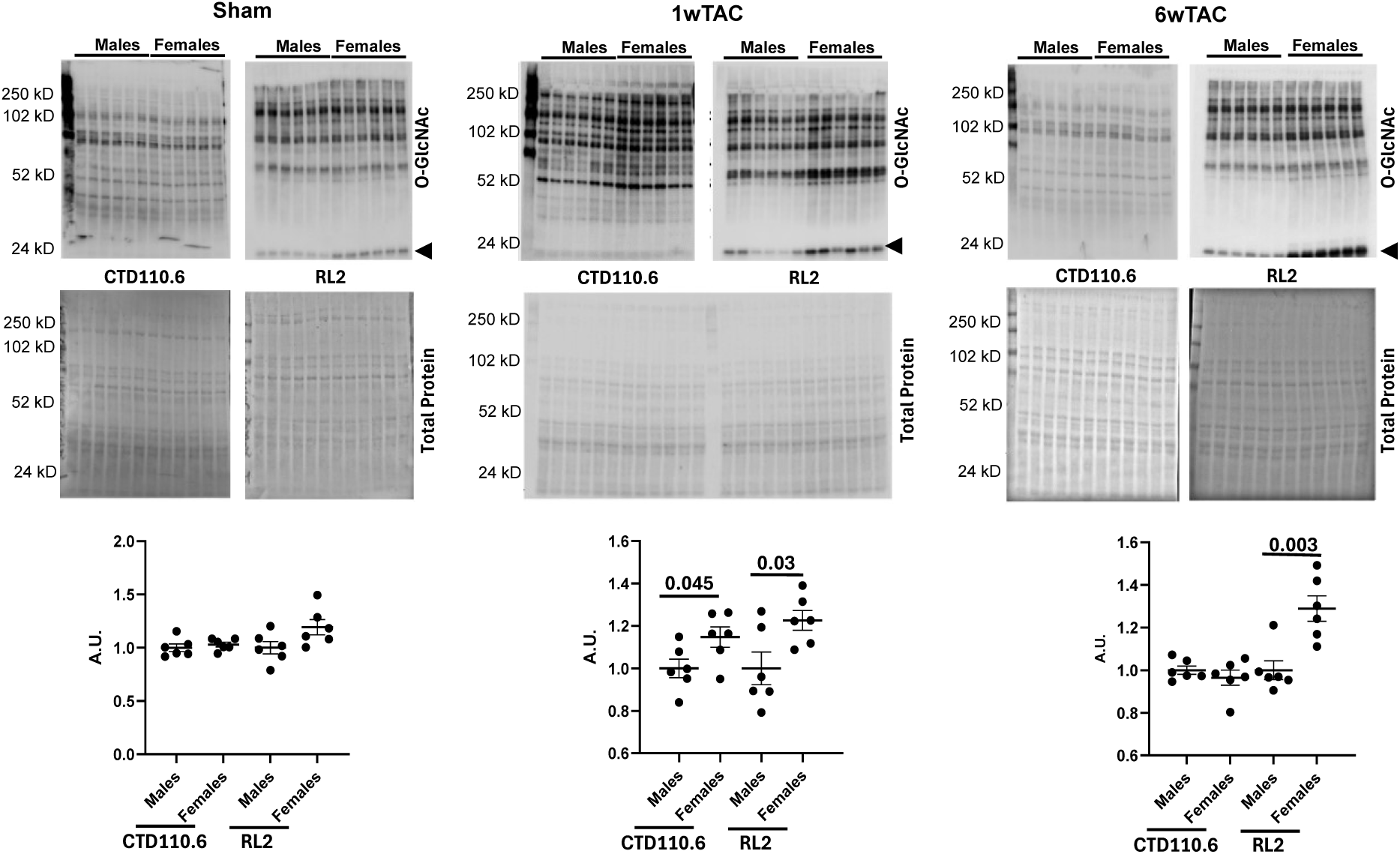
Male to female comparison of global unfractionated cardiac protein O-GlcNAc levels detected by RL2 and CTD110.6. (a) Sham, (b) 1wTAC, and (c) 6wTAC. All values are arbitrary units (A.U.) normalized to total protein levels. Statistical significance was based on unpaired *t*-test (*p* < 0.05). Brackets denote significance between indicated groups. 3, κ (kappa) light chains IgG

### Western Blot Analysis: Subcellular O-GlcNAc Levels

To validate the specificity of our subcellular fractionation protocol, we performed Western blot analysis using antibodies targeting proteins that localize exclusively to distinct cellular compartments. Figure 4 presents representative western blots for each isolated fraction: cytoplasmic, membrane, nuclear, and insoluble pellet. We detected glyceraldehyde-3-phosphate dehydrogenase (GAPDH), a predominantly cytoplasmic protein, exclusively in the cytoplasmic fraction. To evaluate the membrane fraction, we probed for ATP synthase F1 subunit gamma (ATP5C1), a mitochondrial protein, and sarcoplasmic reticulum calcium ATPase 2a (SERCA2a), a membrane-bound calcium transporter. Both proteins appear primarily in the membrane fraction, although overexposure revealed low-level expression of ATP5C1 in the cytoplasmic and pellet fractions (Supplemental Figure S1). For nuclear fraction validation, we used Wilm’s Tumor Associating Protein (WTAP), a ubiquitously expressed nuclear protein. WTAP appeared predominantly in the nuclear fraction, with faint bands in the membrane fraction. Notably, the WTAP antibody also detected a strong band at ∼55 kD in the membrane fraction, whose identity of which remains unknown. We assessed the insoluble pellet fraction using H2AX, a replication-independent histone, and troponin C (TnC), a regulatory protein involved in striated muscle contraction. Both proteins localized exclusively in the insoluble pellet fraction. These compartment-specific marker distributions confirm the effectiveness and fidelity of our fractionation protocol. We subsequently performed fractionated western blot analyses using CTD110.6 and RL2 antibodies across all compartments, stratified by sex (Figures 5-8).

**Figure 4.**
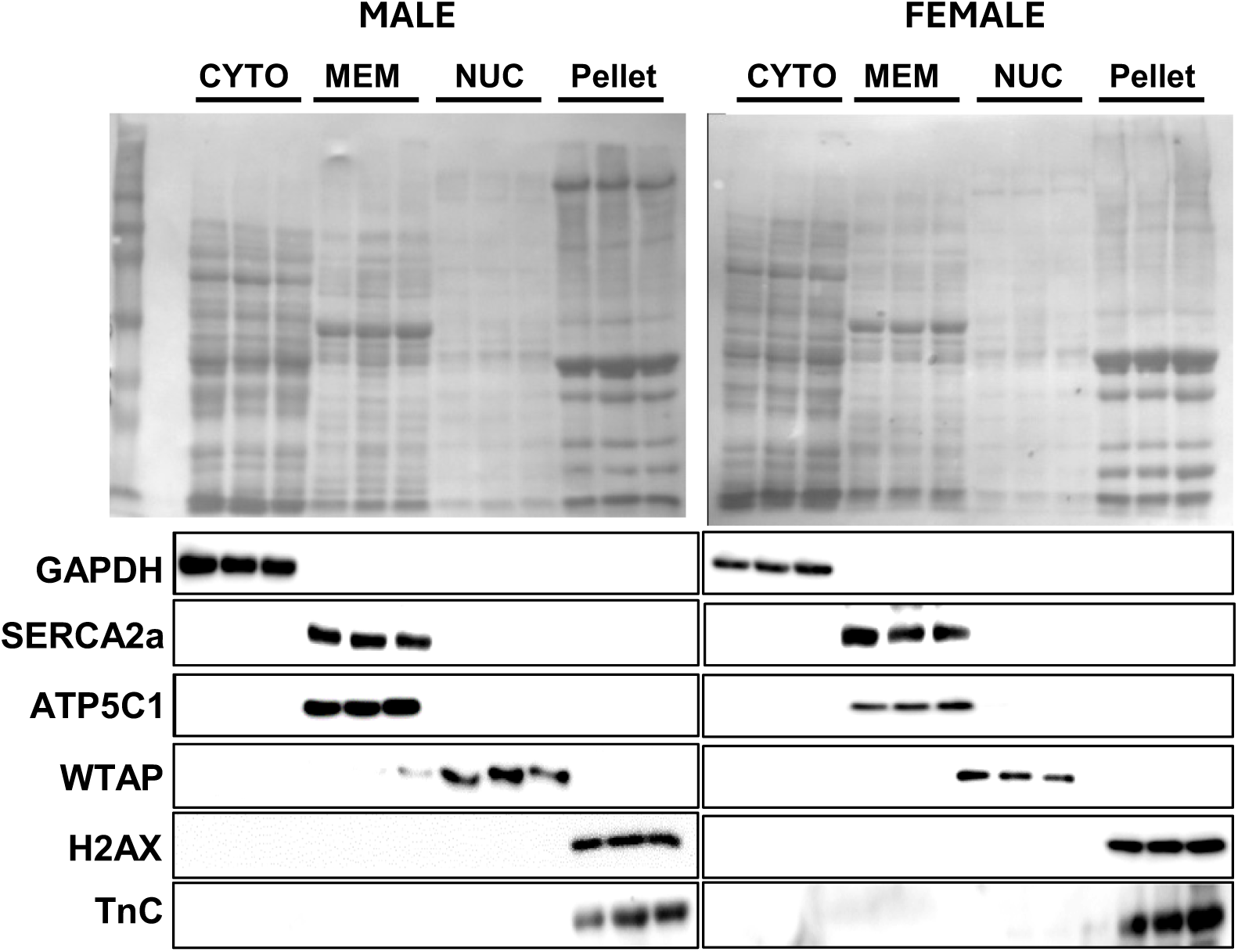
Efficacy of heart tissue subcellular fractionation assessed by western blotting against specific markers. Fractions isolated: cytoplasmic (cyto), marker -GAPDH, glyceraldehyde-3-phosphate dehydrogenase; membrane bound organelles (mem), markers - SERCA, sarcoplasmic reticulum calcium ATPase 2a and ATP5C1, ATP synthase F1 subunit gamma; nuclear (nuc), marker - WTAP, Wilm’s Tumor Associating Protein; and insoluble (pellet) fractions, marker - H2AX, histone H2A.X and TnC, troponin C.

**Figure 5.**
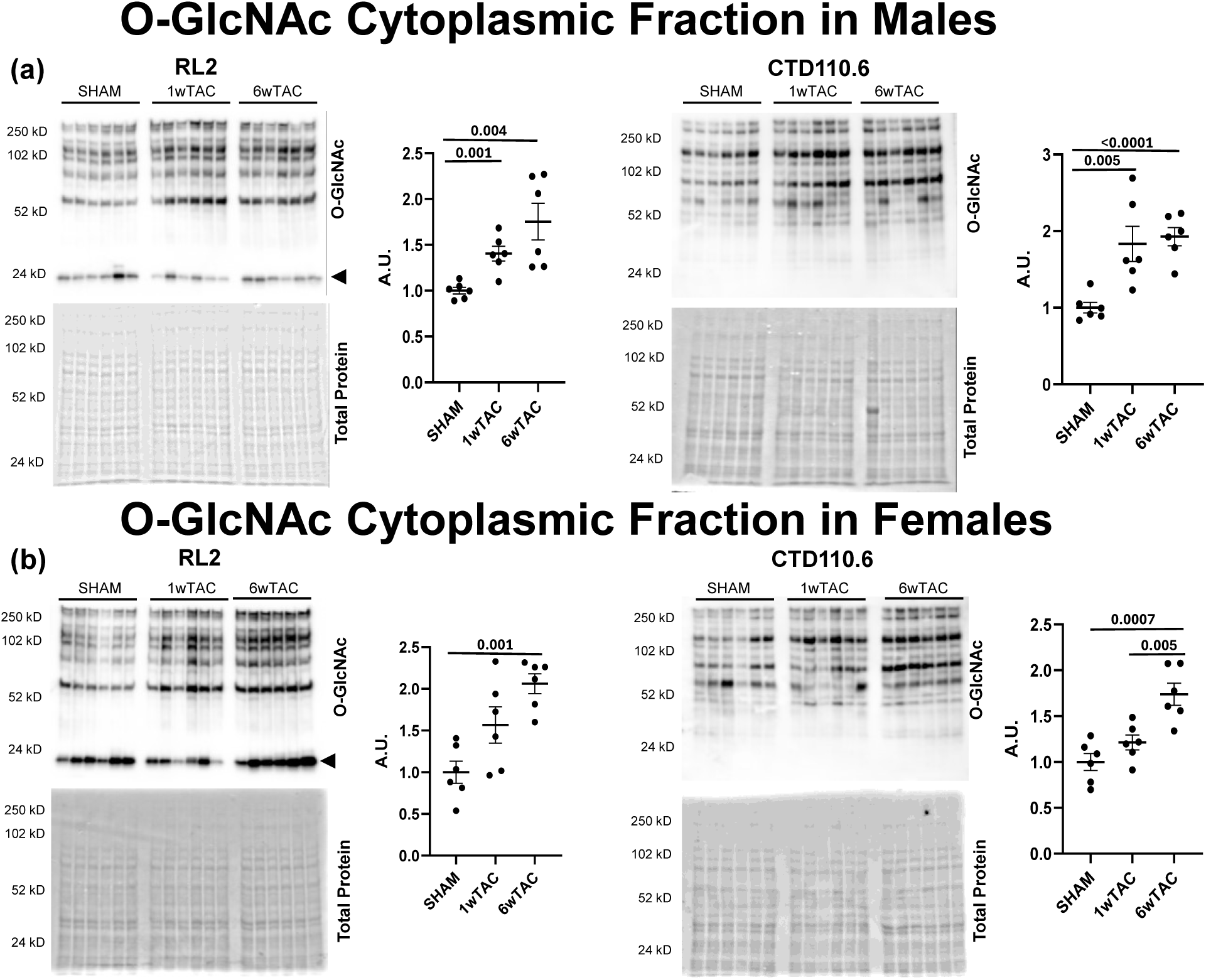
O-GlcNAc levels in the cardiac protein cytoplasmic fraction detected by RL2 and CTD110.6. Males (a) and females (b) comparing Sham, 1wTAC and 6wTAC. All values are arbitrary units (A.U.) normalized to total protein levels. Statistical significance was based on unpaired *t*-test with a Bonferroni correction (*p* < 0.017). Brackets denote significance between indicated groups. 3, κ (kappa) light chains IgG.

In the cytoplasmic fraction (Figure 5), both RL2 (p = 0.001) and CTD110.6 (p = 0.005) showed significantly elevated O-GlcNAc levels in 1wTAC males versus Sham controls. Female 1wTAC had a non-significant trend towards higher cytoplasmic O-GlcNAc levels (p = 0.051) versus Sham control with RL2 but CDT110.6 did not show changes. For the 6wTAC cytoplasmic fraction, both sexes had increased O-GlcNAc levels versus sex-matched Sham controls for both antibodies, RL2 (males, p = 0.004; females, p = 0.001) and CDT110.6 (males, p = <0.0001; females, p = 0.0007). Comparing sex-matched TAC groups, the only significant change was higher cytoplasmic O-GlcNAc levels in female 6wTAC versus 1wTAC with the CTD110.6 antibody (p = 0.005).

In the membrane fraction (Figure 6), we did not detect altered O-GlcNAc levels in 1wTAC compared to Sham for either sex or antibody, although there was a non-significant trend towards higher O-GlcNAc levels in females for the RL2 antibody (p=0.03) only. However, 6wTAC had significantly elevated O-GlcNAc levels over Sham for females detected by both antibodies (RL2, p = 0.003; CTD110.6, p = 0.003), whereas elevated O-GlcNAc levels in males were detected by only the CDT110.6 (p = 0.01. Finally, comparing sex-matched TAC groups, both males and females had higher O-GlcNAc levels in 6wTAC versus 1wTAC with the CTD110.6 antibody only (males, p = 0.001; females, p = 0.002).

**Figure 6.**
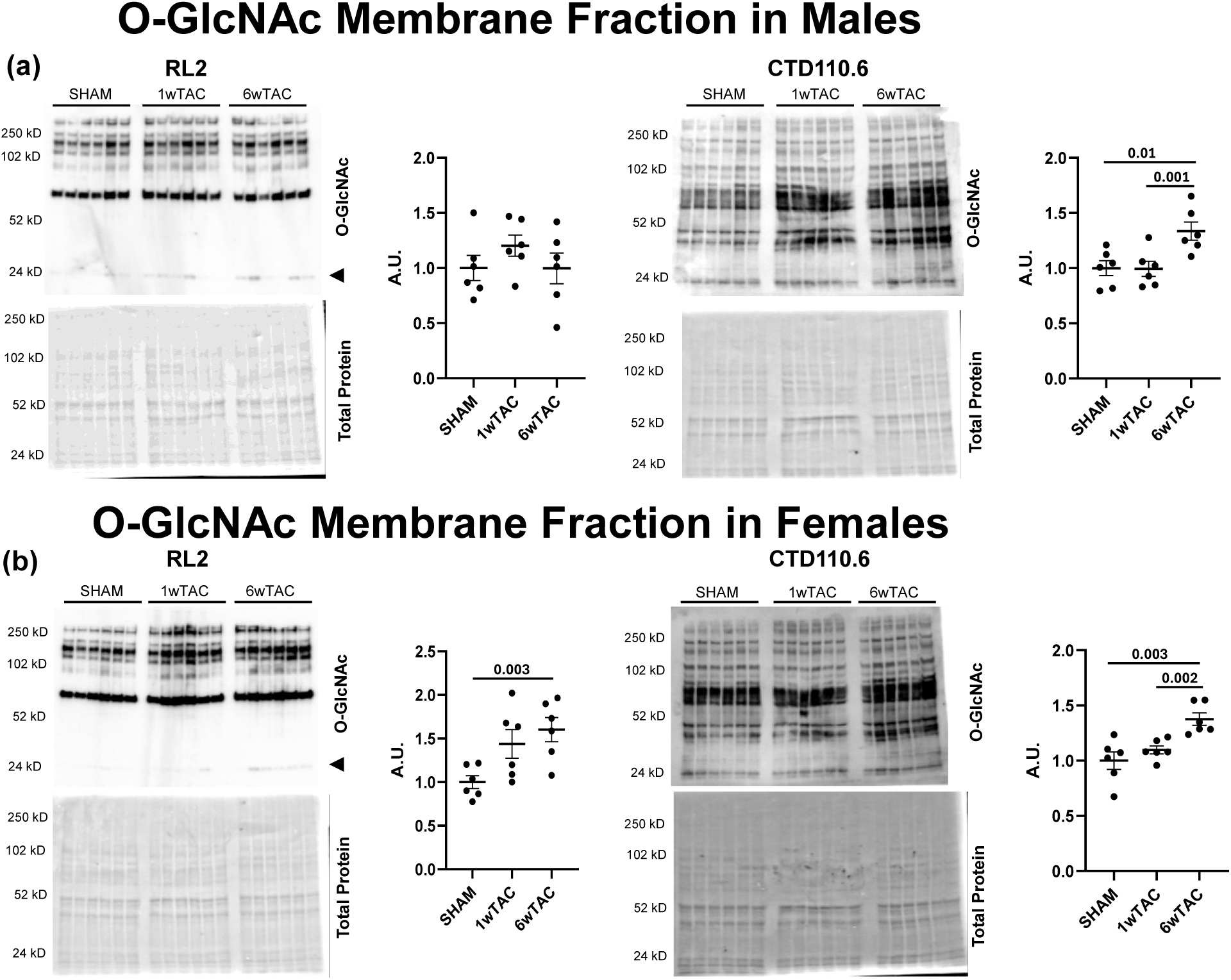
O-GlcNAc levels in the cardiac protein membrane fraction detected by RL2 and CTD110.6. Males (a) and females (b) comparing Sham, 1wTAC and 6wTAC. All values are arbitrary units (A.U.) normalized to total protein levels. Statistical significance was based on unpaired *t*-test with a Bonferroni correction (*p* < 0.017). Brackets denote significance between indicated groups. 3, κ (kappa) light chains IgG.

We did not identify significant changes in O-GlcNAc levels for the nuclear (Figure 7) or contractile fractions (Figure 8) using either antibody.

**Figure 7.**
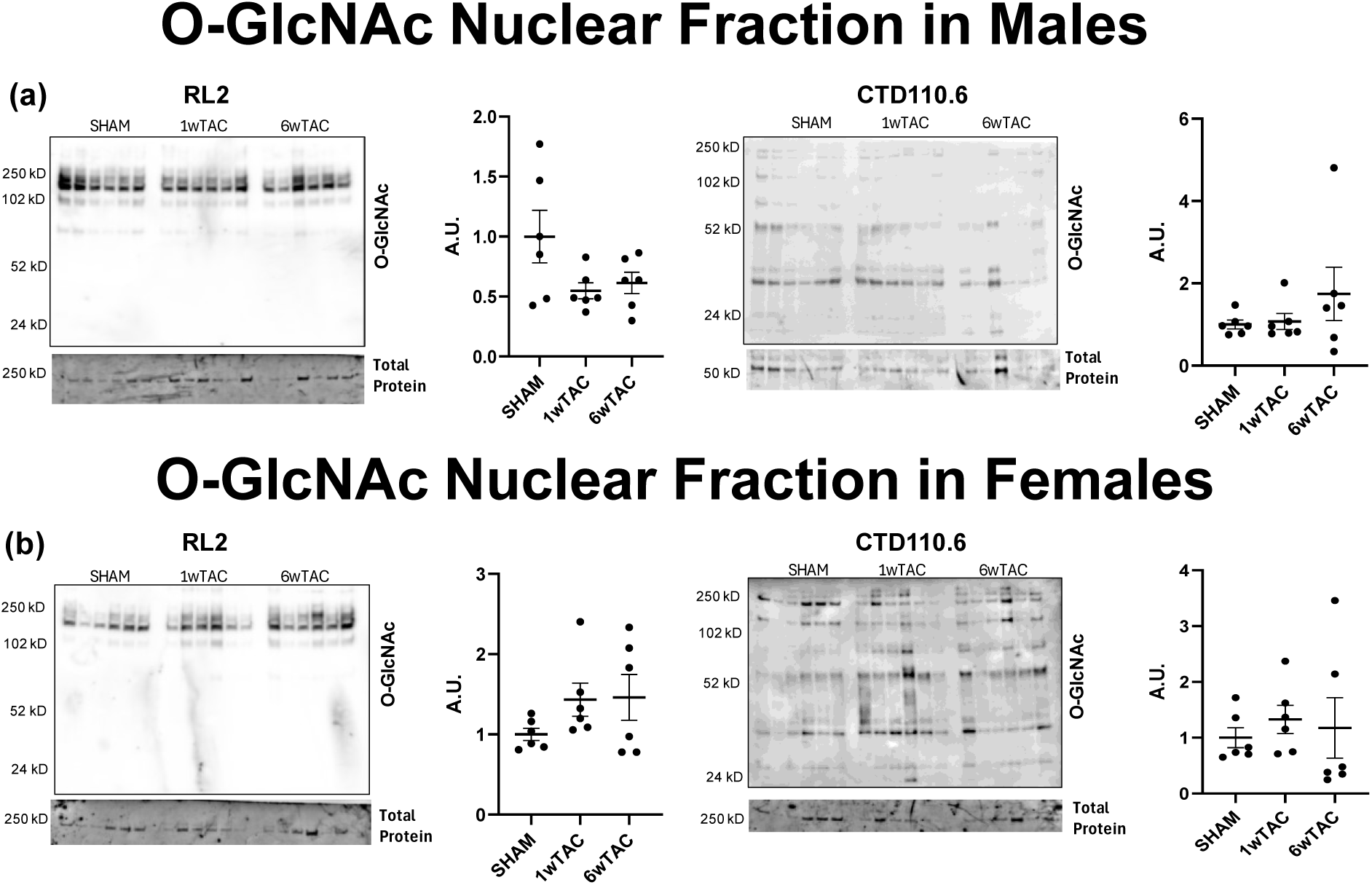
O-GlcNAc levels in the cardiac protein soluble nuclear fraction detected by RL2 and CTD110.6. Males (a) and females (b) comparing Sham, 1wTAC and 6wTAC. All values are arbitrary units (A.U.) normalized to total protein levels. Statistical significance was based on unpaired *t*-test with a Bonferroni correction (*p* < 0.017). Brackets denote significance between indicated groups.

**Figure 8.**
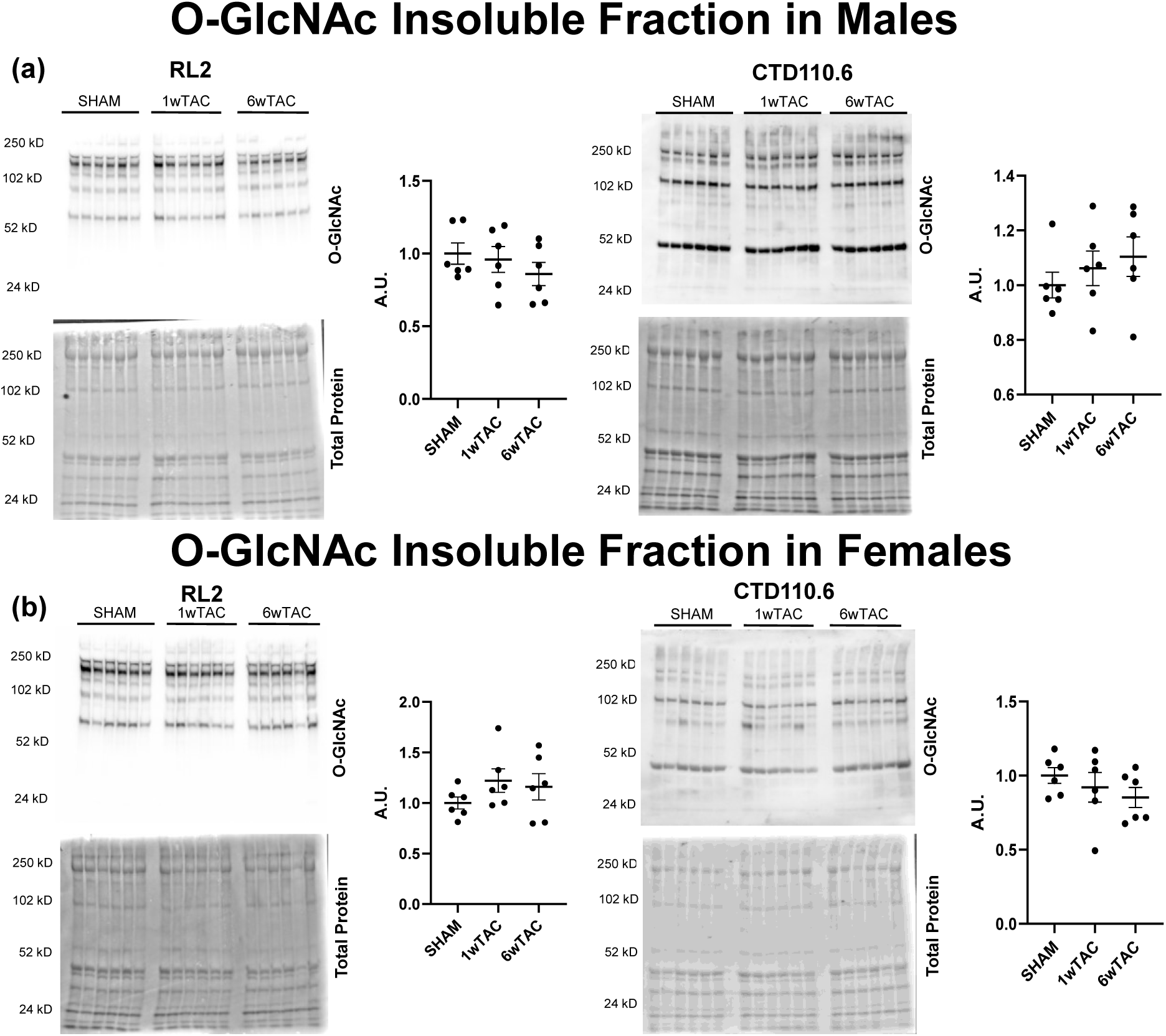
O-GlcNAc levels in the cardiac protein insoluble pellet fraction detected by RL2 and CTD110.6. Males (a) and females (b) comparing Sham, 1wTAC and 6wTAC. All values are arbitrary units (A.U.) normalized to total protein levels. Statistical significance was based on unpaired *t*-test with a Bonferroni correction (*p* < 0.017). Brackets denote significance between indicated groups.

### Western Blot Analysis: Subcellular Regulation of O-GlcNAc by OGT and OGA

Due the differences in O-GlcNAc changes among the compartments, we decided to investigate subcellular regulation of O-GlcNAc by measuring protein levels of the key regulatory proteins OGT and OGA in the same fractions. In the cytoplasmic fraction, OGA protein levels were significantly elevated in 6wTAC compared to Sham in males (p = 0.001) and females (p = 0.0008) (Figure 9a). OGT protein levels were significantly increased in 6wTAC versus Sham for both males (p = 0.0004)and females (p = <0.0001) (Figure 10a).

**Figure 9.**
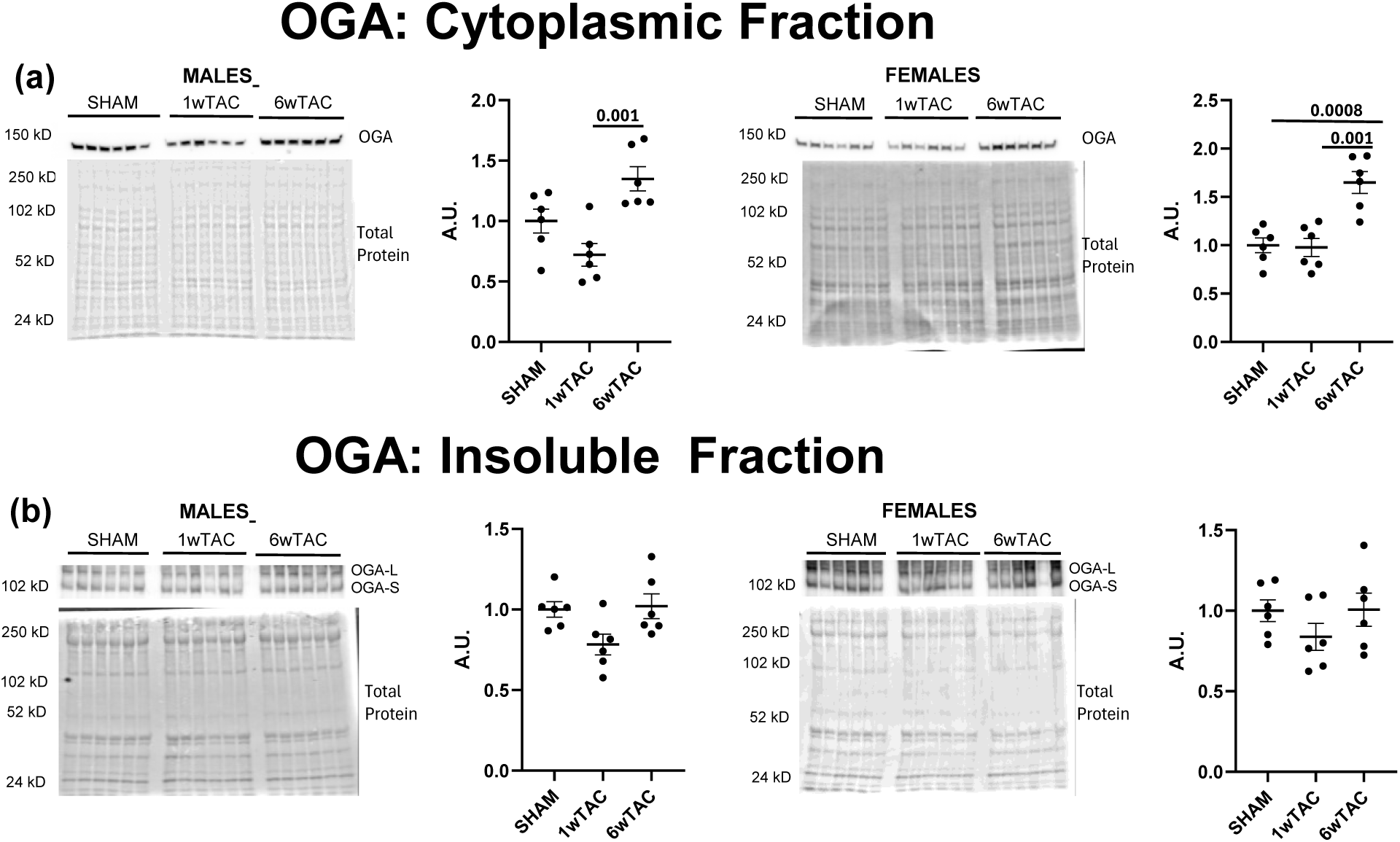
OGA levels in the heart protein cytoplasmic (a) and insoluble pellet (b) fractions for males and female mice comparing Sham, 1wTAC and 6wTAC. All values are arbitrary units (A.U.) normalized to total protein levels. Statistical significance was based on unpaired *t*-test with a Bonferroni correction (*p* < 0.017). Brackets denote significance between indicated groups. (b) Insoluble fraction showed two bands at ∼130 (OGA-long) and ∼100 (OGA-short). Quantification of both bands yielded similar changes in density; therefore, only the graph for OGA-S in shown.

**Figure 10.**
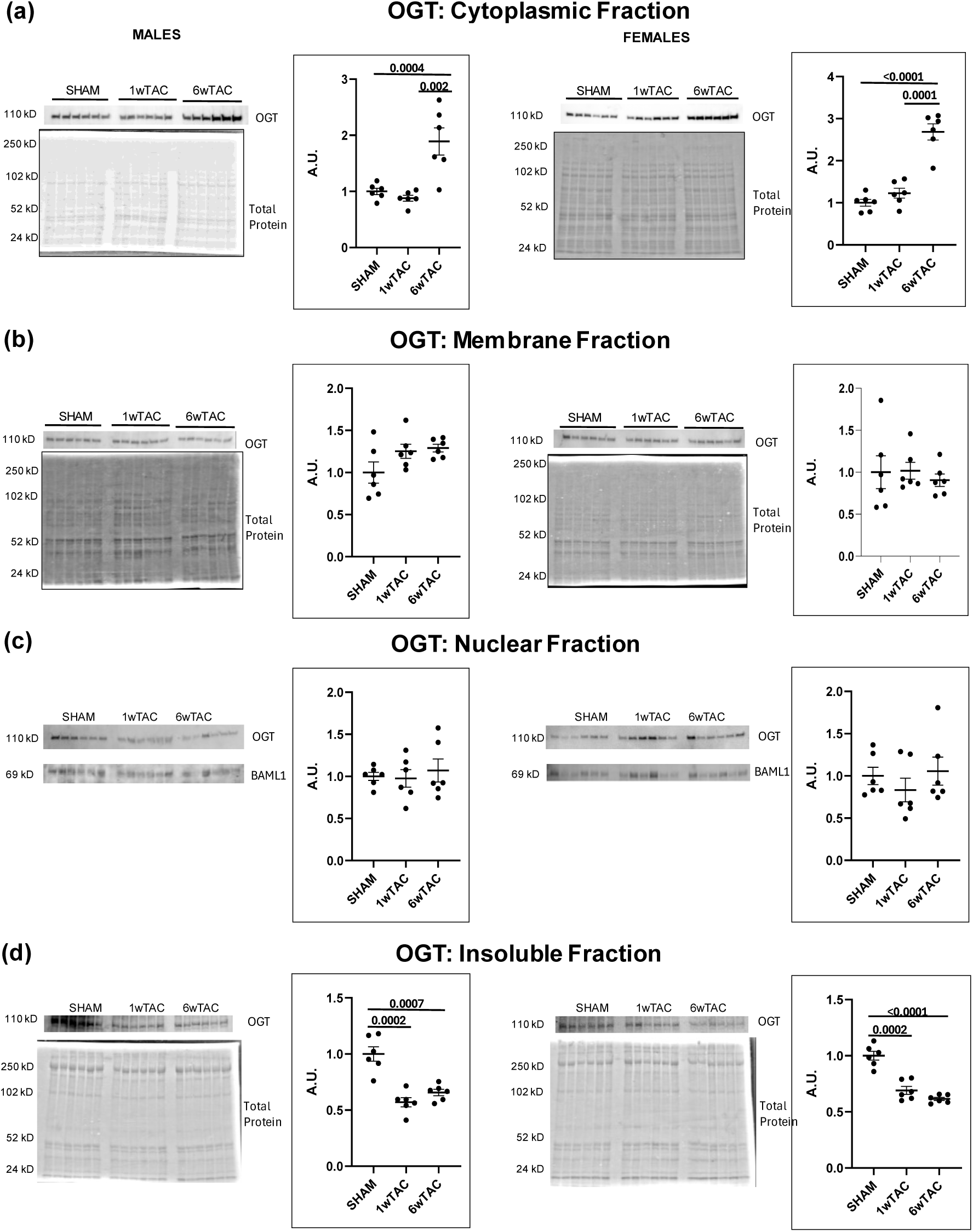
OGT levels in the cardiac protein cytoplasmic (a), membrane (b), nuclear (c) and insoluble pellet (d) fractions for males and female mice comparing Sham, 1wTAC and 6wTAC. All values are arbitrary units (A.U.) normalized to total protein levels.Statistical significance was based on unpaired *t*-test with a Bonferroni correction (*p* < 0.017). Brackets denote significance between indicated groups.

In the Insoluble (pellet) fraction, no detectable changes were observed for OGA across all groups (Figure 9b). OGT levels were significantly decreased in both 1wTAC (males, p = 0.0002; females, p = 0.0002) and 6wTAC (males, p = 0.0007; females, p = <0.0001) groups compared to Sham for both sexes (Figure 10d).

We did not observe significant differences in OGT protein levels in the membrane or nuclear fractions. Unfortunately, we were unable to detect OGA in these fractions under our experimental conditions (Figure 10 b,c).

## DISCUSSION

Protein post-translational modifications by O-GlcNAc play a critical role in regulating tissue and organ function, particularly in response to stressors such as ischemia/reperfusion or pressure overload [4,18]. However, differences in epitope recognition among pan-O-GlcNAc antibodies, coupled with variability in O-GlcNAc immunoblots across laboratories, raise concerns about our field’s ability to reliably detect and characterize dynamic changes in O-GlcNAc signaling. To address these challenges, Zou et al. and Narayanan et al. have recently optimized immunoblotting protocols for the CTD110.6 and RL2 antibodies, respectively, including subcellular fractionation techniques to improve compartment-specific detection [10,11]. Building on these advancements, we demonstrate that adding subcellular fractionation more effectively captures the dynamic O-GlcNAc modifications during POH compared to our previous approach using the RL2 antibody alone on unfractionated samples. Our principal new findings are protein O-GlcNAcylation dysregulation continues from early POH (1wTAC) into chronic POH (6wTAC groups) along with showing differences in O-GlcNAc levels between males and females during POH.

We previously used a similar model to assess early (1wTAC) and chronic (6wTAC) POH for changes in O-GlcNAc levels using the RL2 antibody alone [3]. Our earlier study found global unfractionated O-GlcNAc levels were temporally regulated with increases only in 1wTAC versus Sham controls. Using updated methods and adding the CTD110.6 antibody, we found the same temporal pattern in global unfractionated O-GlcNAc levels with POH. Based on the heart weight/tibial length ratios in our current study, the 1wTAC hearts had not completed hypertrophic growth. Thus, our findings reinforce an important rise in protein O-GlcNAcylation occurring relatively early after TAC and during active hypertrophic growth.

Our new results indicate that O-GlcNAcylation remains an important process in chronic POH. Using cellular fractionation, we observed elevated O-GlcNAc levels in both cytoplasmic and membrane fractions from male and female mice in 6wTAC compared to Sham controls. Our results align with previous findings by Lunde et al., who reported increased O-GlcNAc levels in human patients undergoing surgery for aortic valve stenosis, which is a form of chronic POH in humans [5]. Notably, the O-GlcNAc elevations in their study and our 1wTAC groups were present without employing fractionation. We do not know how detecting O-GlcNAc changes with unfractionated versus fractionated samples manifests on O-GlcNAc protein specificity or impacts specific cellular processes.

While we hypothesize that fewer proteins exhibit elevated O-GlcNAcylation in chronic POH compared to early POH in our model, the extent to which protein specificity is altered remains unclear. For example, we do not know whether the O-GlcNAc elevations in chronic POH represent a smaller subset of same proteins from early POH, fewer but different O-GlcNAcylated proteins, or a combination of these possibilities. This gap in understanding is particularly relevant given that clinical interventions typically occur during the chronic phase of POH. Therefore, elucidating the differences in O-GlcNAc protein specificity between early and chronic POH could be critical for advancing O-GlcNAcylation as a viable therapeutic target.

Although we did not find changes in total O-GlcNAc levels within either the nuclear or pellet fractions during early or chronic POH, we do not interpret these findings as evidence for the absence of significant O-GlcNAc changes on individual proteins within these compartments. Our previous O-GlcNAc proteomics analysis revealed elevated putative O-GlcNAc levels in early POH (also 1wTAC) compared to Sham on proteins localized to these fractions, such as such as heterologous nuclear ribonucleoprotein Q or dynein light chain roadblock type 1 [19]. Taken together, our findings suggest that while cellular fractionation is a useful approach for detecting subtle differences in O-GlcNAc signaling across experimental conditions, it does not preclude the presence of protein-specific modifications even when total compartmental O-GlcNAc levels appear unchanged. This nuance is critical for the design and interpretation of future O-GlcNAc proteomic studies.

While unfractionated global O-GlcNAc levels were significantly elevated in 1wTAC versus Sham for both sexes and antibodies, it is notable that the fractionation differences were primarily in 6wTAC. We do not have an explanation or hypothesis for this mild divergence. It would be informative for other investigators to compare unfractionated and fractionated results to see whether they observe a similar phenomenon.

Although our focus was on determining O-GlcNAc levels, we also performed a limited screen of the key O-GlcNAc regulatory proteins OGA and OGT within the fractions. We previously assessed unfractionated OGT and OGA during POH and found changes in these regulatory proteins which would be consistent with augmenting O-GlcNAcylation with early POH [3]. Specifically, we found that the increase in global O-GlcNAc levels during early POH were associated with a fall in OGA levels in males, whereas females had both a fall in OGA and an increase in OGT [3]. However, we did not find a similar pattern with fractionation in the current study. The higher cytoplasmic O-GlcNAc levels in 6wTAC versus Sham for both sexes was associated with increases for both OGT and OGA. We interpret these cytoplasmic results to suggests that OGT activity increases to a greater extent than OGA to promote O-GlcNAcylation in this fraction. In the membrane fraction, we did not find protein level changes that would lead to increased O-GlcNAcylation which suggests enzyme activities are affected post-translationally. An alternative possibility for higher O-GlcNAc levels in the cytoplasm and membrane fractions is that O-GlcNAcylation promoted localization of outside proteins to these fractions. Overall, our findings show the limitations with using protein quantification alone to understand O-GlcNAc regulation especially within cellular compartments.

### Female-Male differences in O-GlcNAc

Our current study identified greater total O-GlcNAc elevations in females during POH. Specifically, total O-GlcNAc levels were increased in females over males at 1wTAC for both antibodies and at 6wTAC for RL2 only. This finding differs from our prior study using the RL2 antibody alone where we found similar O-GlcNAc levels between the sexes in 1wTAC. Upon further review, we now recognize that the quality of the prior immunoblot was suboptimal because the banding pattern is not wholly consistent with the other RL2 immunoblots in that study or our current study. Thus, our new finding illustrates the challenges with O-GlcNAc western blot variability even within our lab.

We did not observe sex-based differences in total O-GlcNAc levels in Sham hearts using either antibody. This finding diverges from Narayanan et al., who reported higher O-GlcNAc levels in female hearts compared to males using RL2, both at baseline (control hearts) and under ischemic conditions [20]. While sex differences in O-GlcNAc responses to stress appear consistent across the studies, the reasons for different results in control hearts are unknown. There are minor methodological variations between our laboratories which may contribute and warrant further investigation [10,11]. For example, Narayanan et al. [20] used 9 M urea lysis buffer and has gifted custom RL2 antibody whereas we use RIPA lysis buffer and a commercial RL2 antibody.

Collectively, our findings reinforce the presence of sex-specific differences in cardiac O-GlcNAcylation. In addition to the work by Narayanan et al. [20], we previously demonstrated that hypothyroidism elevates total O-GlcNAc levels in aged female hearts but not in males. We also identified sex-dependent expression patterns of O-GlcNAc regulatory proteins in the heart under thyroid hormone perturbations or POH [3,9]. Narayanan et al. attributed increased O-GlcNAc signaling in female hearts to enhanced activity of O-GlcNAc transferase (OGT) [20], the sole enzyme responsible for O-GlcNAc addition, but we did not evaluate OGT activity in this study. Future investigations should account for sex differences in cardiac O-GlcNAcylation when examining cardiac O-GlcNAcylation.

As with our discussion on early versus chronic POH, it is essential to determine how sex-based differences in O-GlcNAc levels under stress conditions such as POH or ischemia influence protein-specific modifications and downstream cellular functions. Future studies should also explore whether O-GlcNAcylation contributes to sex disparities in cardiovascular outcomes.

### Technical Insights for Future O-GlcNAc Studies in the Heart

The relative utility of the CTD110.6 versus RL2 antibodies for assessing cardiac O-GlcNAc levels remains an area of active discussion. CTD110.6 is known to recognize a wider range of proteins but may exhibit limited reactivity with cardiac contractile proteins [11]. In contrast, RL2 binds cardiac contractile proteins effectively but shows reduced sensitivity toward low molecular weight proteins [11]. In our study, overall O-GlcNAc trends were largely consistent between the two antibodies, with a few exceptions. Specifically, RL2 uniquely detected elevated O-GlcNAc levels in females compared to males in 6wTAC, and in female 1wTAC versus Sham hearts within the membrane fraction (trend only). Conversely, CTD110.6 alone identified differences in the membrane fraction for males and females (1wTAC versus 6wTAC only). Our results suggest that either antibody can be suitable for cardiac studies, if investigators follow the key methodological recommendations outlined by Zou et al. [10] for CTD110.6 and Narayanan et al. [11] for RL2 and validate consistent banding patterns under their specific immunoblotting practices.

With improved molecular weight coverage on our immunoblots, we a priori hypothesized that there could be O-GlcNAc differences between Sham and 6wTAC at a few specific protein bands that become obscured when measuring global O-GlcNAc levels. However, we identified only one area with reproducible band-level difference, an increase in 6wTAC hearts compared to Sham for RL2 O-GlcNAc signal at 102 kD in females. Given that cell fractionation revealed O-GlcNAc differences between Sham and 6wTAC in both sexes, we propose that fractionation may offer a more sensitive approach for detecting subtle changes in O-GlcNAcylation across experimental conditions than analyzing specific molecular weight bands, at least within cardiac tissue. This raises important considerations regarding the utility of fractionation in O-GlcNAc immunoblot workflows.

Our data suggest that fractionation is not necessary when standard immunoblots reveal O-GlcNAc differences between experimental groups. However, in studies involving chronic stressors where conventional whole-organ preparations fail to detect changes in O-GlcNAc levels, fractionation may enhance detection sensitivity. To further develop this workflow, we encourage investigators to incorporate fractionation into their O-GlcNAc immunoblot protocols and report the outcomes in their publications.

### Limitations

First, our POH hearts demonstrated stable systolic function suggesting that they had compensated hypertrophy. We do not know whether our findings apply to POH with decompensated left ventricular function and heart failure findings. Second, we incorporated many, but not all, of the O-GlcNAc immunoblot steps from Zou et al. [10] and Narayanan et al. [11]. Nevertheless, we optimized methods in our lab to achieve consistent immunoblot results with a similar range in molecular weights for the banding. Third, there are multiple methods for fractionation prior to immunoblots. We did not test whether specific fractionation methods are better or worse in O-GlcNAc immunoblots. Fourth, we only utilized the two most common antibodies for cardiac O-GlcNAc western blots. We do not know whether other O-GlcNAc antibodies or non-immunoblot techniques like Click-it chemistry would be better at detecting subtle O-GlcNAc changes in chronic POH without fractionation. Finally, we used C57/Bl6J mice, and we do not know whether similar O-GlcNAc changes occur with other mouse strains or species. Thus, it would be informative to repeat similar studies in other strains and species.

### Conclusions

In summary, our study highlights the dynamic regulation of cardiac protein O-GlcNAcylation in response to pressure overload hypertrophy (POH), with distinct temporal and sex-specific patterns. Using updated immunoblotting techniques and cellular fractionation, we demonstrate that O-GlcNAc modifications are increased starting during early hypertrophic growth (1wTAC) and persisting into chronic POH (6wTAC). Further, female mice exhibited greater O-GlcNAc elevations than males during POH, underscoring the importance of considering sex as a biological variable in future O-GlcNAc studies. Our findings also emphasize the need for careful methodological consistency, especially around immunoblot variability, because they can influence detection outcomes. Ultimately, exploring protein-specific O-GlcNAcylation and its impact on cellular functions will be critical for advancing its role as a therapeutic target in cardiac disease.

## Availability of data and materials

All data generated or analyzed during this study are available from the corresponding author on reasonable request.

## Authors’ contributions

DL, AO, and WZ conceived and designed research; DL, AO, and WZ. performed experiments; DL, AO, and WZ. analyzed data; DL, AO, and WZ interpreted results; DL and AI prepared figured and drafted manuscript; DL and AO edited and revised manuscript. DL, AO, and WZ approved the final version of the manuscript.

## Acknowledgements

Not applicable.

## Supplemental Data

Supplemental figures 1-4 show the uncropped western blots.

