## Supplemental Tables for "Detection of Cardiac O-GlcNAcylation via Subcellular Fractionation and Dual Antibody Analysis in Pressure Overload Cardiac Hypertrophy"

SUPPLEMENTAL FIGURES

Figure S1. Complete images of fractionation confirmation blots.

Figure S2. Uncropped blots of the membranes used in detection of OGA.

Figure S3. Uncropped blots of the membranes used in detection of OGT.

Figure S4. Uncropped blots of the membranes used in nuclear fraction O-GlcNAc detection.

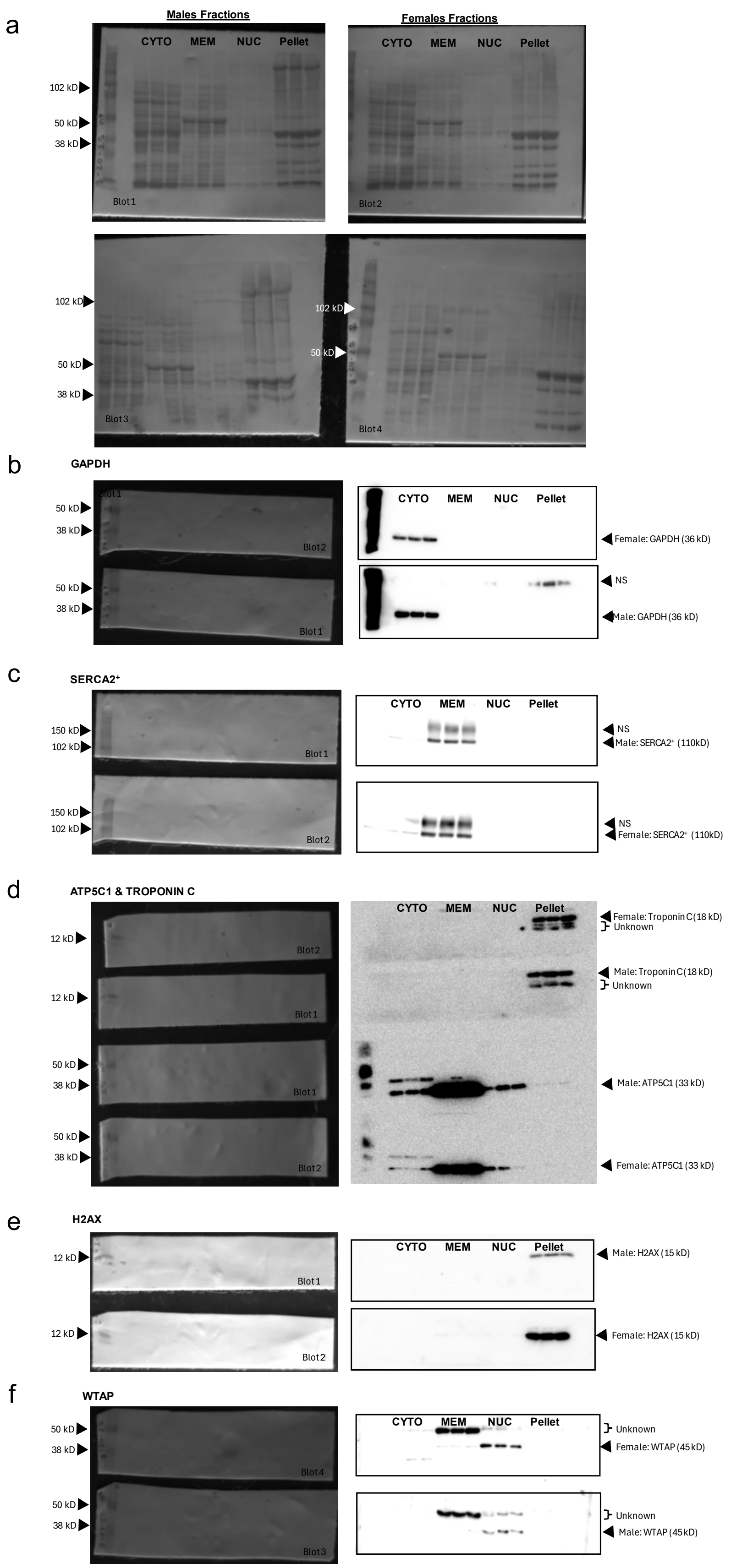

a
Males Fractions
Females Fractions
CYTO MEM NUC Pellet
102 kD
50 kD
38 kD
Blot1
Blot2
Blot3
Blot4
b
GAPDH
Female: GAPDH (36 kD)
NS
Male: GAPDH (36 kD)
c
SERCA2+
150 kD
102 kD
NS
Male: SERCA2+ (110kD)
Female: SERCA2+ (110kD)
d
ATP5C1 & TROPONIN C
12 kD
Female: Troponin C (18 kD)
Unknown
Male: Troponin C (18 kD)
Male: ATP5C1 (33 kD)
Female: ATP5C1 (33 kD)
e
H2AX
Male: H2AX (15 kD)
Female: H2AX (15 kD)
f
WTAP
Unknown
Female: WTAP (45 kD)
Male: WTAP (45 kD)

Supplemental Figure 1. Fractionation check. (a) depicts protein stain of blots containing fractionated protein (n = 3). (b) Cytoplasmic fraction confirmation with GAPDH antibody. (c) Membrane fraction confirmation with mitochondrial SERCA2$^{+}$ antibody. (d) Membrane fraction check with mitochondrial ATP5C1 antibody and insoluble fraction check with contractile protein troponin C antibody. (e) Insoluble fraction check with chromatin H2AX antibody. (f) Nuclear fraction check with WTAP antibody.

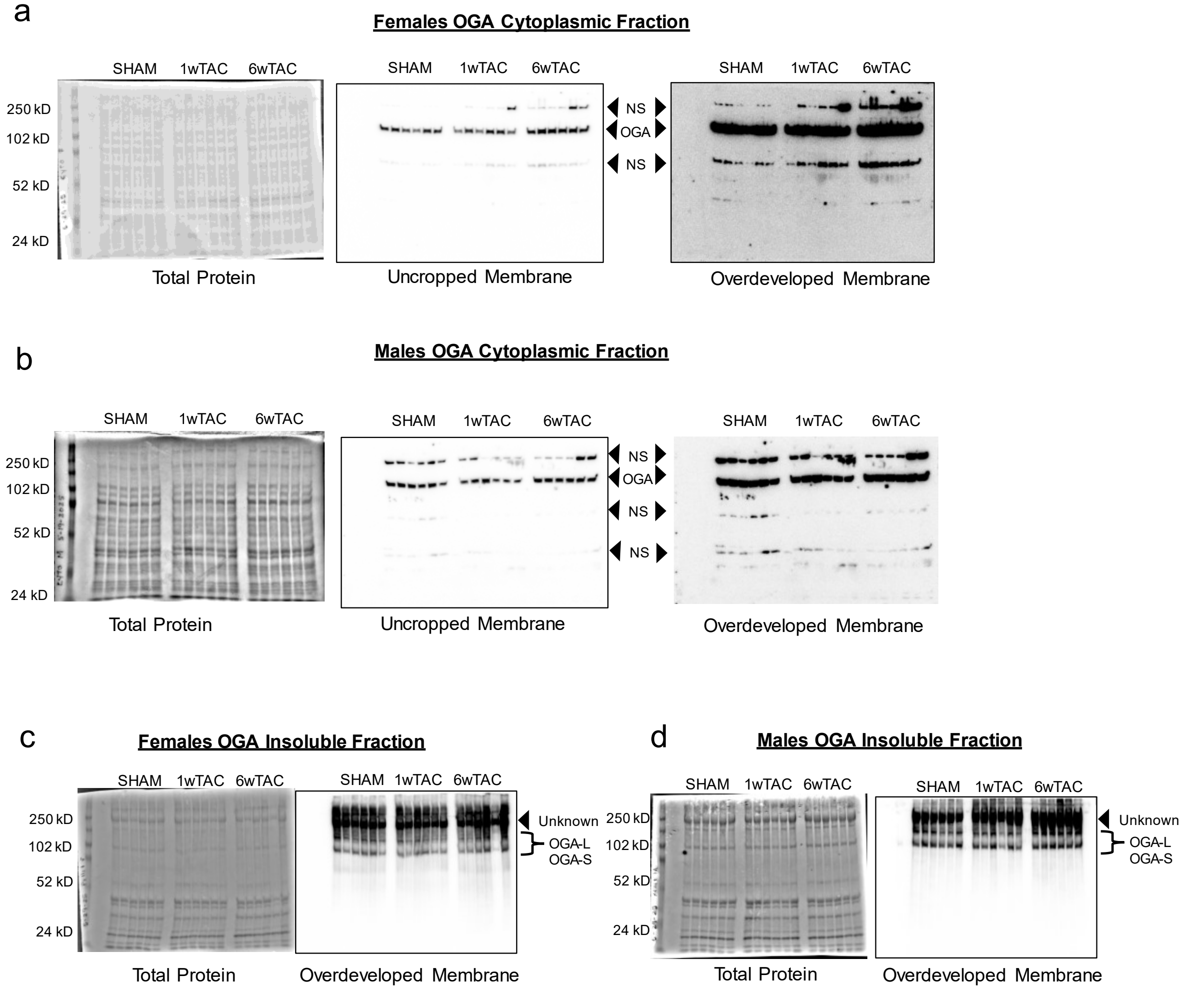


Supplemental Figure 2. OGA levels in the heart protein cytoplasmic fraction (a) females (b) males and insoluble fraction (c) females (d) males. Cytoplasmic fraction: OGA band at 130 kD observed. Additionally, faint nonspecific bands are observed in the cytoplasmic fraction at ~250, 76 and 40 kD (a and b). Insoluble fraction: An unknown band is detected at ~250 kD with OGA bands observed at ~130 and ~100 kD in the insoluble fractions (c and d).

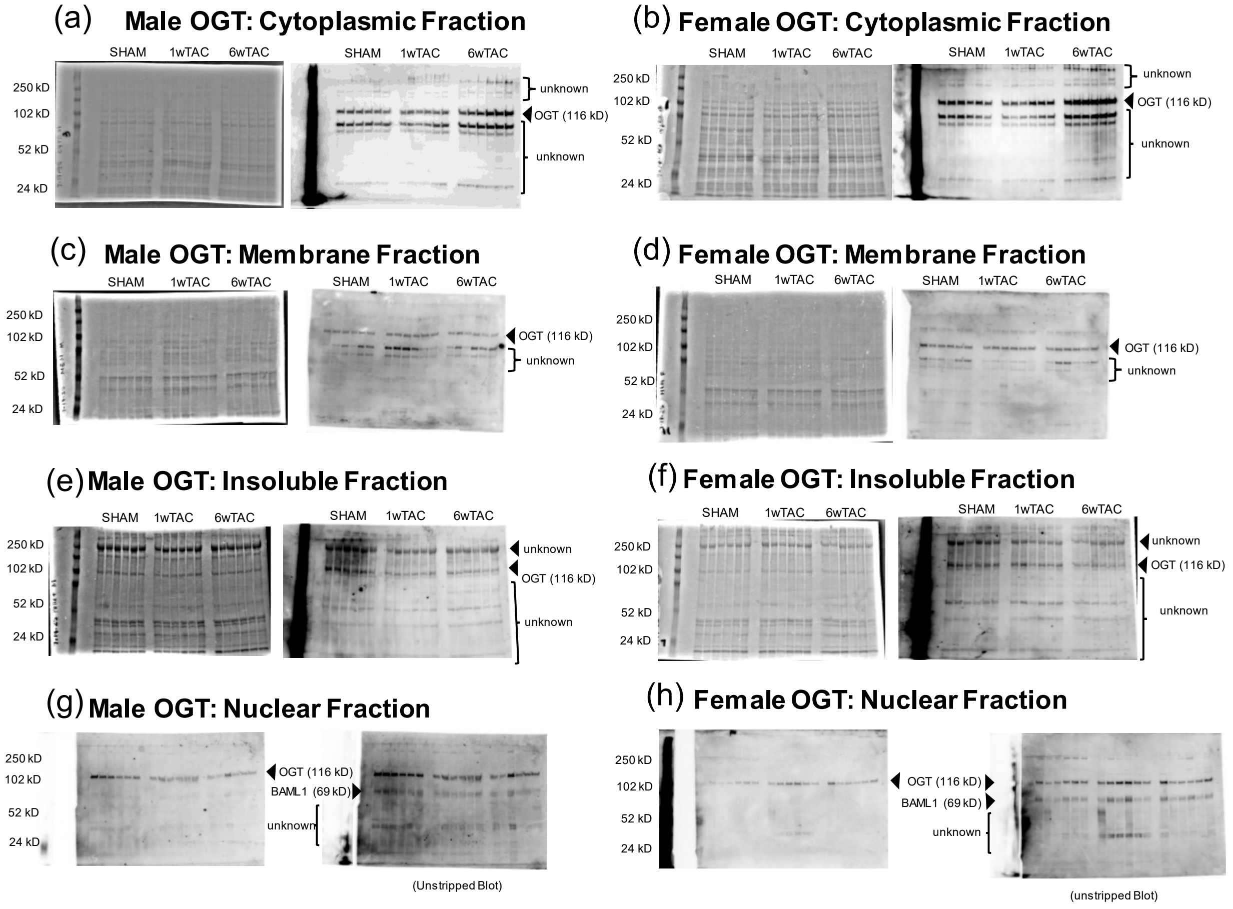


Supplemental Figure 3. OGT levels in the heart protein cytoplasmic fraction (a) males (b) females, membrane fraction (c) males and (d) females, insoluble fraction (e) males (f) females, nuclear fraction (g) males (h) females. insoluble fraction (c) females (d) males. Cytoplasmic fraction (a and b): OGT band at 116 kD. Faint high molecular weight bands between 250 and 150 kD. Two bands observed at ~76 kD region (possible isoforms, not quantified). Membrane fraction (c and d): OGT band at 116 kD. Two bands observed at ~76 kD region (possible isoforms, not quantified). Insoluble fraction (e and f): OGT band at 116 kD. high molecular weight bands at ~250 kD and lower molecular weights below 50 kD. Nuclear fraction (g and h): Unstripped OGT membrane probed with BAML1 antibody shows the OGT band at 116 kD and the BAML1 band at ~70 kD. Nonspecific lower molecular weights are also observed.

### (a) O-GlcNAc Nuclear Fraction in Males

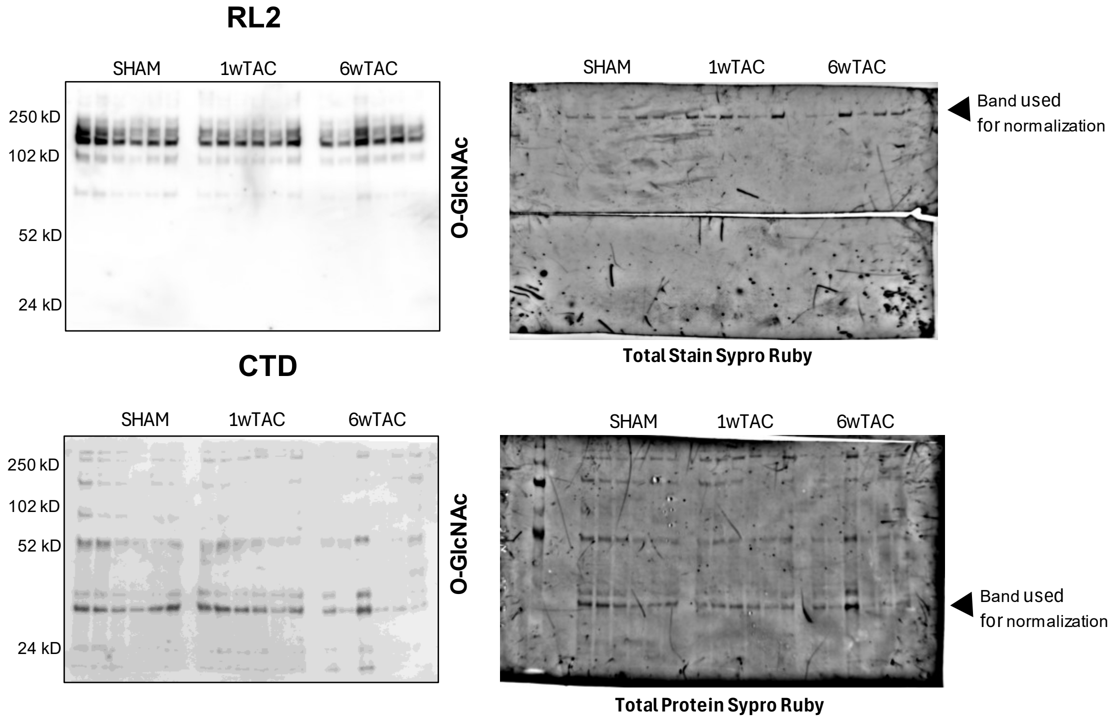


### (b) O-GlcNAc Nuclear Fraction in Females

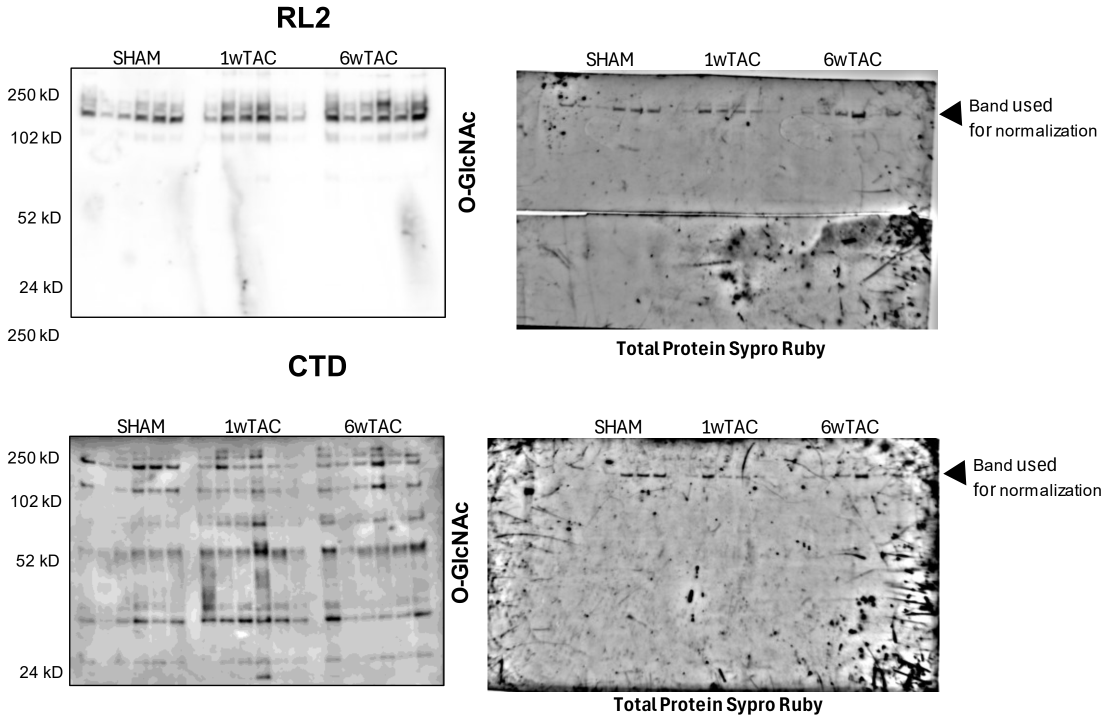

Supplemental Figure 4. O-GlcNAc levels in the cardiac protein soluble nuclear fraction detected by RL2 and CTD. Males (a) and females (b). Normalized to a protein band detected using Sypro Ruby. Loading of nuclear protein at 5 mg made detection by staining problematic. Blots had been cut for use with other antibodies before being stained with Sypro Ruby.
